# Sonic Hedgehog is not a limb morphogen but acts as a trigger to specify all digits

**DOI:** 10.1101/2020.05.28.122119

**Authors:** Jianjian Zhu, Rashmi Patel, Anna Trofka, Brian D. Harfe, Susan Mackem

## Abstract

Limb patterning by Sonic hedgehog (Shh) has served as a paradigm of “morphogen” function, acting either by graded spatial or temporal signal integration. Yet how Shh instructs distinct digit identities remains controversial. Here, we bypassed the requirement for Shh in cell survival during limb bud outgrowth and demonstrate that a transient, early Shh pulse is both necessary and sufficient for normal limb development. Our analysis of Shh-responding cells shows signaling acts at only short-range during this time window and that Shh patterns digits indirectly, via a relay mechanism, rather than by direct spatial or temporal signal integration. Using a genetic assay for relay signaling, we unexpectedly discovered Shh also specifies digit 1 (the thumb; previously thought to be exclusively Shh-independent) indirectly, thus implicating Shh in a unique regulatory hierarchy for digit 1 evolutionary adaptations, such as opposable thumbs. This study illuminates Shh as a trigger for a relay network that becomes rapidly self-sustaining, with mechanistic relevance for limb development, regeneration, and evolution.

## Introduction

Shh has been considered a prototypical vertebrate morphogen, with the limb and neural tube serving as mainstays to elucidate its function. In contrast to neural tube, anterior-posterior (A-P) limb patterning yields different digit identities from a common set of cell types and tissues, with distinct skeletal morphologies that are not based in cell fate changes per se, but arise as an emergent property. Understanding how Shh specifies limb skeletal pattern is central to the problem of structural morphogenesis and informative to regenerative medicine. Shh is secreted by posterior limb bud mesoderm cells defined functionally as the “zone of polarizing activity” (ZPA) and regulates the specification of distinct A-P digits 2-5 (d2-d5; index to pinky)(reviewed in ^1^), excepting d1 (the thumb), which has been thought to form Shh-independently^2^. Patterning and growth are coupled and during its 2-day expression span in limb bud (Fig. 1A), Shh controls both digit type and number.

**Figure 1.**
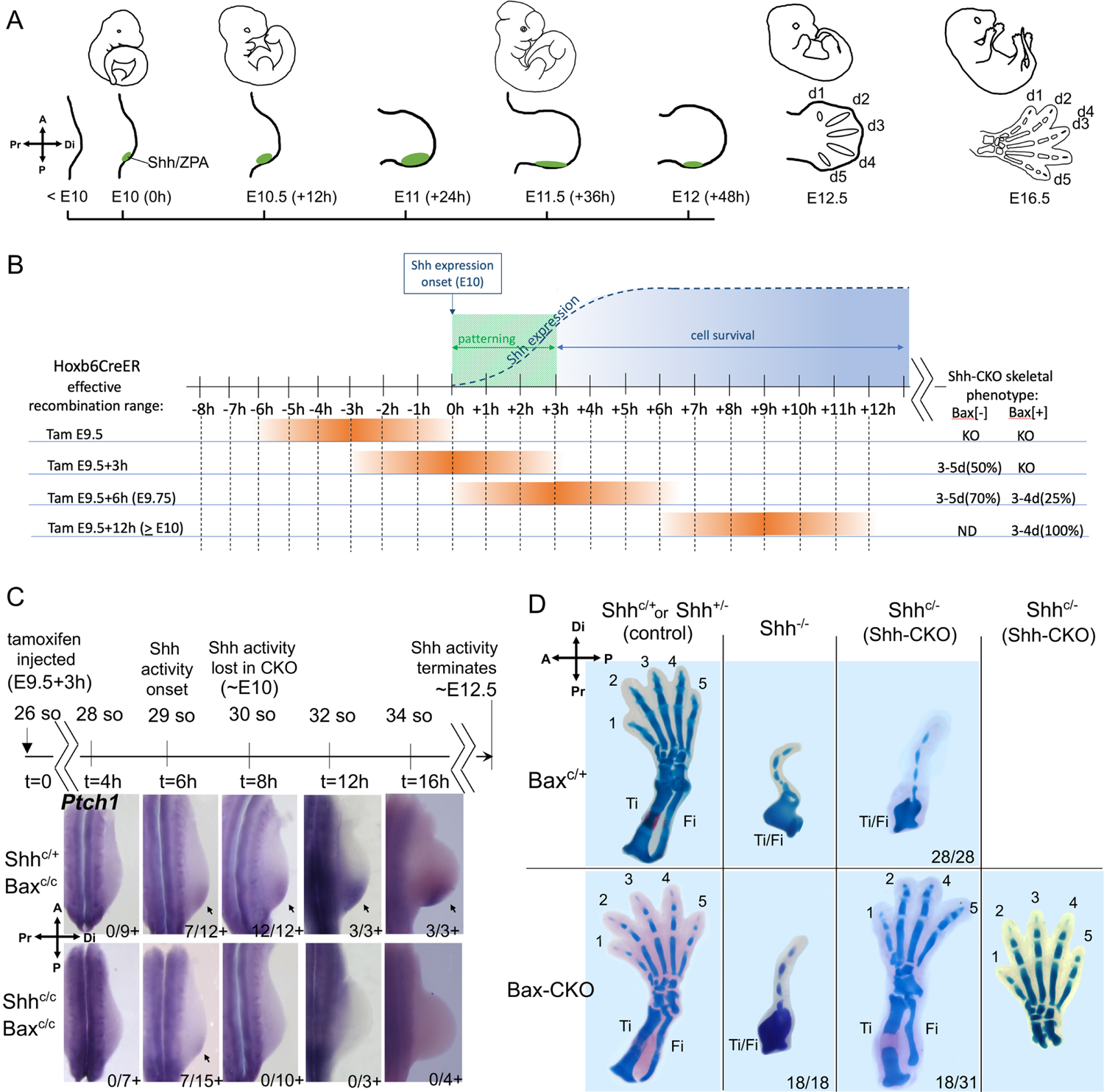
A transient Shh activity pulse specifies all digits with enforced cell survival. **(A)** Shh expression/functional timelines in wildtype mouse hindlimb buds (Shh/ZPA in green). Limb axis orientation indicated by compasses (shown at left) in all figures. **(B).** Summary of recombination timelines (orange) for early limb bud tamoxifen-induced deletion of Shh at different treatment times and subsequent digit outcomes (at right; KO, Shh null phenotype; ND not done), shown relative to timing of Shh expression onset. At E9.5 treatment, deletion precedes Shh expression and all limbs have a Shh null phenotype. By E9.75 and later, deletion occurs after the Shh role in maintaining cell survival has already commenced and both Bax[+] (apoptosis competent) and Bax[-] (cell survival enforced) embryos display some rescue of digit formation. >E10 data summarized from Zhu et al (2008). See also Table S1. **(C)** Shh activity assayed by *Ptch1* RNA expression (arrows) at times after tamoxifen, as indicated on timeline. Shh activity was first detected at 6hr after tamoxifen injection in a subset of both control *Shh*^+/C^;*Bax*-CKO (7/12+), and in *Shh*-CKO;*Bax*-CKO (7/15+) embryos, and became robust by 8hr in control (12/12+), but was absent in all *Shh*-CKO;*Bax*-CKO embryos (0/10+) from t=8hrs on. so, somite. **(D)** Skeletal staining (E16.5) following *Bax/Bak* and *Shh* removal by tamoxifen (treated at E9.5+3h, as in **C**). In hindlimbs with *Bax/Bak* present (Bax^c/+^) all *Shh*-CKO embryos (28/28) have *Shh* null phenotype. In hindlimbs with *Bax/Bak* absent (Bax-CKO), about half of the *Shh*-CKO embryos (18/31) have 3-5 normal digits (5-digit phenotype shown in right-most panel) and normal zeugopod bones (tibia, Ti; fibula, Fi), but all *Shh* null embryos (*Shh*^-/-^; 18/18) with *Bax/Bak* removed still retain the null mutant phenotype. Related to Figure S1 and Table S1.

Yet, despite high functional conservation across tetrapods and intense investigation for over two decades, the mechanism by which Shh patterns digits remains controversial. Very disparate models have been proposed in different species with similar limb gene regulatory networks^1, 3–6^. Morphogen-based models of Shh function derive from chick studies showing that changes in concentration or duration can alter both digit identity and numbers, with posterior identities requiring higher concentrations or longer exposures^3, 6, 7^. Lineage tracing of ZPA-descendants in both chick and mouse have revealed that d4 and d5 arise from the descendants of Shh-expressing ZPA cells^4, 8^, leading to the proposal that temporal integration of short-range Shh, rather than graded long-range signaling, specifies digit identities. While non-ZPA cells are displaced further away from ZPA signals during outgrowth, Shh-expressing d4,d5 progenitors receive the longest exposure^4^. But the normal coupling of Shh roles in digit identity (patterning) and number (growth) poses a challenge to tease apart the individual requirements for each. Comparing the effects of Shh inhibition with that of cell cycle blockade to reduce digit number in chick^3^, led to the proposal that temporal signal integration acts to progressively “promote” digit territories to more posterior identity. Yet genetic lineage tracing indicated that Shh-expressing ZPA cells become refractory to Shh-response over time^9^, seemingly at odds with temporal integration models to achieve posterior identity. Using a conditional *Shh*-mutant we previously found *Shh* was required for only a limited interval (∼8-10 hrs) to specify normal digits; later *Shh* removal caused digit loss in an order that reflected the normal temporal order in which digits arise, and not their A-P position. Accordingly, the remaining digits that formed were morphologically normal and not transformed to an anterior identity (*5*). These results suggested a biphasic model in which Shh is required early for patterning, but over a more extended time to promote progenitor survival and expansion, potentially restricting Shh-morphogen action to a more limited time frame.

Here, we use a genetic strategy to uncouple the Shh requirement in digit patterning from expansion/cell survival and test how Shh acts to specify digit pattern. We demonstrate that a transient (∼2hr) burst of Shh activity, during which the Shh-response is limited to d4-d5 progenitors suffices to specify all digits normally, implying that non-ZPA derived digits are patterned through an indirect, relay mechanism and that Shh acts neither as a spatial morphogen nor via temporal integration. A genetic assay to test for Shh-induced relay signals unexpectedly reveals that d1 specification is in fact Shh-dependent, requiring an indirect downstream relay signal and highlighting a role for Shh in the evolutionary emergence of a unique, polarized thumb (digit 1).

## Results

### A transient Shh pulse restores normal limb development when cell survival is enforced

To examine the early role of Shh selectively, we used a genetic strategy to uncouple the Shh requirement in digit patterning from that in outgrowth. We asked if enforcing cell survival would bypass the late-phase Shh function, and restore any digit formation. The pro-apoptotic *Bax*/*Bak* genes^10^ were deleted in *Shh* conditional mutant limb buds exposed to a very transient, early Shh pulse (using *Shh*^C/C^;Hoxb6CreER, hereafter referred to as *Shh*-CKO, and *Bax*^C/C^;*Bak*^-/-^ alleles, referred to as *Bax*-CKO; see Table 1 for complete list of all crosses and genotypes used). Our analysis focused on hindlimb, where Hoxb6CreER drives complete and robust recombination even at very early stages before limb bud initiation (∼E9; see Fig 1A)^11^. Since this very early recombination will delete Shh prior to its expression onset, while deletion at too late a time will produce only mild digit loss phenotypes even despite reduced cell survival^5^, the tamoxifen timing to induce Cre and delete *Shh* was optimized (to E9.5+3hrs, Figure 1B and Table S1) so that 100% of *Shh*-CKO sibling embryos that retained one wild-type *Bax* allele (cell survival not enforced) were invariably *Shh* germ-line null (*Shh*^-/-^) in hindlimb skeletal phenotype (28/28), providing a clear baseline to assess the effects of bypassing *Shh* late function. This tamoxifen treatment time (E9.5+3h, Fig. 1B, Table S1) resulted in rescued digit formation in about half of embryos with enforced cell survival (Fig. 1D; discussed below) and precedes the normal onset of *Shh* expression by about 9 hours^5, 12^. Earlier deletion (at E9.5) invariably produced a null phenotype regardless of whether enforced cell survival was present (Table S1), consistent with previous work showing that E9.5 treatment deletes *Shh* efficiently prior to expression onset^5^. At later deletion times (E9.5+6h), some partial rescue of digit formation occurred even in a portion of embryos without enforcement of cell survival (∼25%, see Table S1).

**Table 1.**
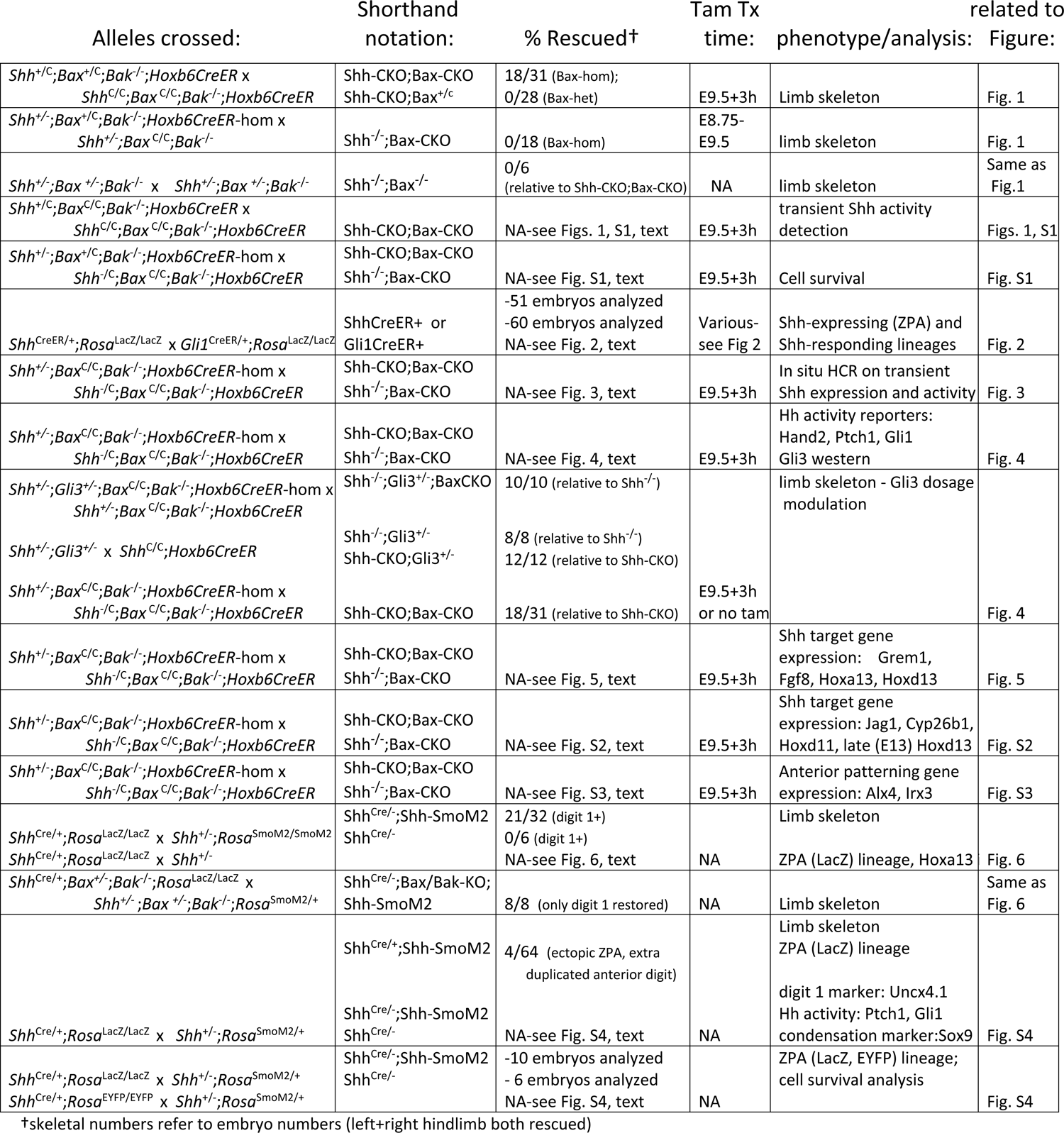
List of genetic crosses, embryo numbers analyzed, and outcomes for experiments related to each figure.

To determine the total duration of Shh signaling activity after an optimized tamoxifen treatment time (of E9.5+3h), Shh response was assayed by detecting the direct Shh target RNAs (*Gli1*, *Ptch1*)^13–15^ at 2-hour (∼one somite) intervals post-tamoxifen treatment (Figs. 1C, S1A). Direct target detection is a highly sensitive and earlier read-out for Shh function than is the presence of Shh RNA itself^5, 12^, and at later stages also serves as a sensitive indicator for any mosaic recombination, since residual Shh-expressing cells proliferate over time and would become readily apparent at later stages. Analysis of Shh-response at 2-hour intervals after tamoxifen detected a transient 2-3 hour window of Shh activity in about half of the *Shh-*CKO embryos (7/15 *Ptch1+*, 4/10 *Gli1+*, Figs. 1C, S1A). No activity was detected in the remaining half, presumably because recombination occurred prior to *Shh* expression onset in those embryos. Notably, transient *Shh* activity was also detected in only half of control (*Shh*^+/-^) embryos at this stage/time, in agreement with the early tamoxifen deletion timing relative to *Shh* expression onset (Fig. 1B,C, S1A; see also Table S1). Within two hours later (+1 somite), when Shh activity (*Ptch1*, *Gli1* expression) is detected in all control embryos, Shh activity has ceased in all *Shh*-CKO embryos (Fig 1B,C; Fig S1A) and remains absent subsequently. Analysis of *Shh-*CKO;*Bax-*CKO embryos at multiple later stages also failed to detect the late emergence of any direct target *Gli1* or *Ptch1* expression (see below).

Cell survival was completely restored in 100% of *Shh* mutant limb buds with *Bax-*CKO (20/20, Fig. S1B), but notably *Bax*-CKO alone had no effects on limb skeletal patterning in *Shh*^+/C^;*Bax*-CKO sibling controls (Fig. 1D). In *Shh*-CKO;*Bax-*CKO embryos, blocking cell death restored formation of from 3 to 5 digits with normal morphology in about half the embryos (18/31); the remaining 50% retained the *Shh*^-/-^ null mutant limb phenotype (13/31; Fig. 1D), correlating well with the fraction of embryos that did not display any transient Shh activity earlier (Figs. 1C, S1A; Table S1) and which subsequently have a *Shh* null skeletal phenotype (Fig 1D, Table S1). In *Shh-*CKO;*Bax*-CKO embryos with rescued limbs (18/31), normal long bone morphology (tibia/fibula; zeugopod) was also restored (see also Fig. 2C), and normal AP polarity was clearly evident in both the long bones and digits, including distinctive d1 and d5 identities. In contrast, all (100%) of sibling *Shh*-CKO embryos retaining one functional *Bax* allele (*Shh*-CKO;*Bax*^+/C^) had levels of apoptosis very similar to the Shh^-/-^; *Bax*^+/C^ (Fig S1B) and later displayed a *Shh* null limb skeletal phenotype with a single dysmorphic digit and malformed zeugopod (28/28, Fig. 1D).

**Figure 2.**
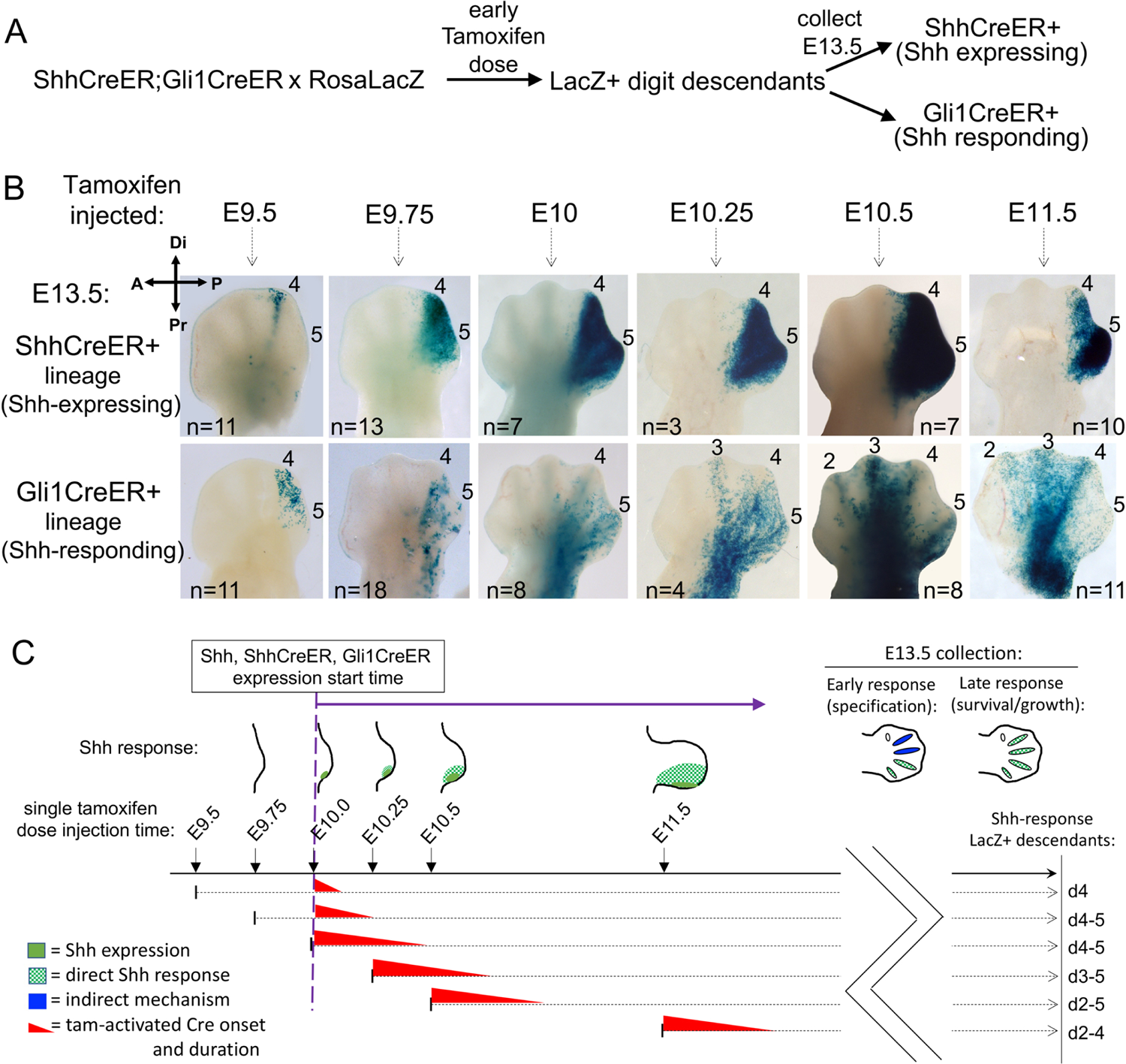
Lineage tracing of *Shh* response in normal limb buds at the time of *Shh* onset. **(A)** Diagram of strategy to compare Shh-expressing and Shh-responding lineages in sibling embryos so that developmental ages and tamoxifen timing conditions are identical. **(B)** Distribution of RosaLacZ reporter+ descendants after a single tamoxifen dose injected at the stage indicated, and collected at E13.5 to visualize digit rays. After E9.5 or E9.75 tamoxifen (time window as in Fig. 1B), direct Shh-response is limited to digit 4, 5 territory (ZPA domain; Shh-expressing descendants). Long-range response in digit 2-3 territory is evident by E10.5. n, numbers analyzed for each injection time. **(C)** Summary of data in (**B**) showing overlap of tamoxifen activity time window (red wedge) with *Shh* expression/response (solid/hatched green in hindlimb bud schematics) and listing digit contributions of Shh-responsive descendants (to right) for each tamoxifen injection time. Effective tamoxifen duration was estimated at ∼12 hours^61^ (see also from Figs. 1B,C, S1A). Early-responding cells (tamoxifen at E9.5 or E9.75) only contribute to d4, d5 (hatched green in E13.5 hindlimb), indicating d2, d3 are specified by an indirect mechanism (blue). Late-responding cells (tamoxifen after E10.25), during phase when Shh acts to maintain cell survival, contribute to d2 – d5 (hatched green in E13.5 hindlimb).

These data indicate that a transient early Shh pulse (∼2-3h) suffices to specify digit progenitors, but not to maintain cell survival. We next asked if a short Shh pulse is even necessary when cell survival is enforced, using a non-conditional *Shh* null mutant completely devoid of all Shh activity (*Shh*^-/-^;*Bax-*CKO; Fig. 1D). Although cell death was completely blocked in *Shh*^-/-^;*Bax-*CKO limbs (9/9, Fig. S1B), the enforced cell survival failed to rescue any digit or normal long bone formation (0/18, Fig. 1D), even when a non-conditional germ-line *Bax/Bak* mutant (*Shh*^-/-^;*Bax*^-/-^;*Bak*^-/-^) was used to ensure complete *Bax/Bak* inactivation in the *Shh*^-/-^ (0/6 skeletal rescue; Table 1). These results indicate that an early, transient Shh pulse (of 2-3hr) is both necessary and sufficient for normal digit and long bone formation, if the role of *Shh* in maintaining cell survival is bypassed by *Bax/Bak* removal. Consequently, later stage sustained Shh signaling acts mainly to support cell survival and limb bud expansion.

### Rescued digits in the early Shh-CKO with enforced cell survival arise from cells that did not respond directly to Shh

The short 2-3h Shh activity window required for normal limb development when cell survival is enforced is inconsistent with temporal signal integration models, but could be compatible with transient activity of a spatially graded morphogen. To assess whether Shh acts as a long-range morphogen across the limb bud during this early time window in the normal developing limb bud, we used lineage tracing with *Gli1*^CreER/+^ ^4, 9^ to track cells that had responded to Shh at early times immediately after *Shh* expression onset. We also compared Shh-response (Gli1CreER activity) with the spatial extent of the Shh-producing ZPA region at the same times (*Shh*^CreER/+^)^4^ as a guide to assessing the extent of long-range signaling. Lineage tracing crosses included both *Shh*^CreER/+^ and *Gli1*^CreER/+^ knock-in alleles to genetically mark cells (RosaLacZ reporter activation) in sibling embryos from the same litter that expressed either *Shh*^CreER/+^ or *Gli1*^CreER/+^ alone, ensuring that Shh production and Shh response were compared at identical embryonic ages and tamoxifen exposure times (Fig. 2A). A single tamoxifen dose was given at closely spaced early times spanning *Shh* (and *ShhCreER*) expression onset (Fig. 2B,C) and limb buds were collected at E13.5, after all digit rays have formed, so that the descendant (LacZ+) cell contributions to different digits can be easily scored (Fig. 2A,B).

Genetic LacZ reporter marking revealed that Shh acts only very short-range at early times after expression onset. For tamoxifen treatment prior to E10.25, Shh response (Gli1CreER activation) was confined to cells that later give rise to d4, d5 (Fig. 2B). Long-range signaling was not detected until much later tamoxifen-induction times, initially extending toward d3 (E10.25) and later also including d2 territories (by E10.5, Fig. 2B). Yet tamoxifen treatment at a much earlier stage (at E9.5+3h, Fig. 1), provides a transient Shh pulse that is both necessary and sufficient to specify all 5 digits if cell survival is enforced. During this time interval, the lineage tracing shows that Shh acts only short-range. Shh response marked by Gli1CreER activity is limited to the Shh-producing ZPA region and only directly impacts only progenitors that later give rise to d4, d5. Consequently, other “Shh-dependent” digit progenitors must respond indirectly to Shh, indicating an indirect relay mechanism (Fig. 2C-blue). Notably however, at later stages Shh does act as a long-range signal (by E10.5, Fig. 2B and 2C-green-hatched), in the context of its ongoing later role in maintaining cell survival.

Is the same time frame for short and long range Shh signaling also maintained in the *Shh-CKO;Bax-CKO* mutant? The requirement for a robust, early Cre driver to achieve efficient, rapid deletion in the Shh-CKO;Bax-CKO precludes genetic lineage analysis using Cre lines to track Shh response in the mutant context. Gli1LacZ is an alternative response reporter over short time frames (see below), but cannot be used to assess the digit contributions of early responding progenitors because early-induced LacZ protein does not perdure over the 3-day time interval between transient *Shh* induction and digit ray formation (∼E10 to E13.5). To determine if the extent of Shh response relative to early *Shh* expression is similar to wildtype in the *Shh*-CKO during the short period of Shh activity following tamoxifen treatment (data in Fig. 1), we used hybridization chain reaction (HCR)^16^ to provide a very sensitive and high resolution spatial detection of *Shh* RNA expression and response (*Gli1* RNA expression) simultaneously, by multiplex labeling in the same limb bud to accurately compare spatial distributions. Note that the *Shh* probes used detect recombined as well as functional *Shh* transcripts, and that midline hindgut *Ihh* signaling, preserved in the *Shh*^-/-^,^17^ activates some proximal axial response in the urogenital-cloacal region^18, 19^ (see *Shh* and *Gli1* RNAs in Fig. 3 *Shh*^-/-^ panels). Using HCR simultaneous detection (Fig. 3), *Gli1*+ response is very similar in its A-P extent to *Shh* and is short range in both the control and *Shh*-CKO mutant during the transient pulse when response is detected in the mutant limb bud (Fig. 3, 29 somite panels, arrows). This confirms that, as in the wildtype, Shh activity/response in the *Shh*-CKO;*Bax*-CKO mutant with rescued digit formation is limited to short-range pathway activation during the transient *Shh* pulse, in the territory that will give rise to digit 4-5.

**Figure 3.**
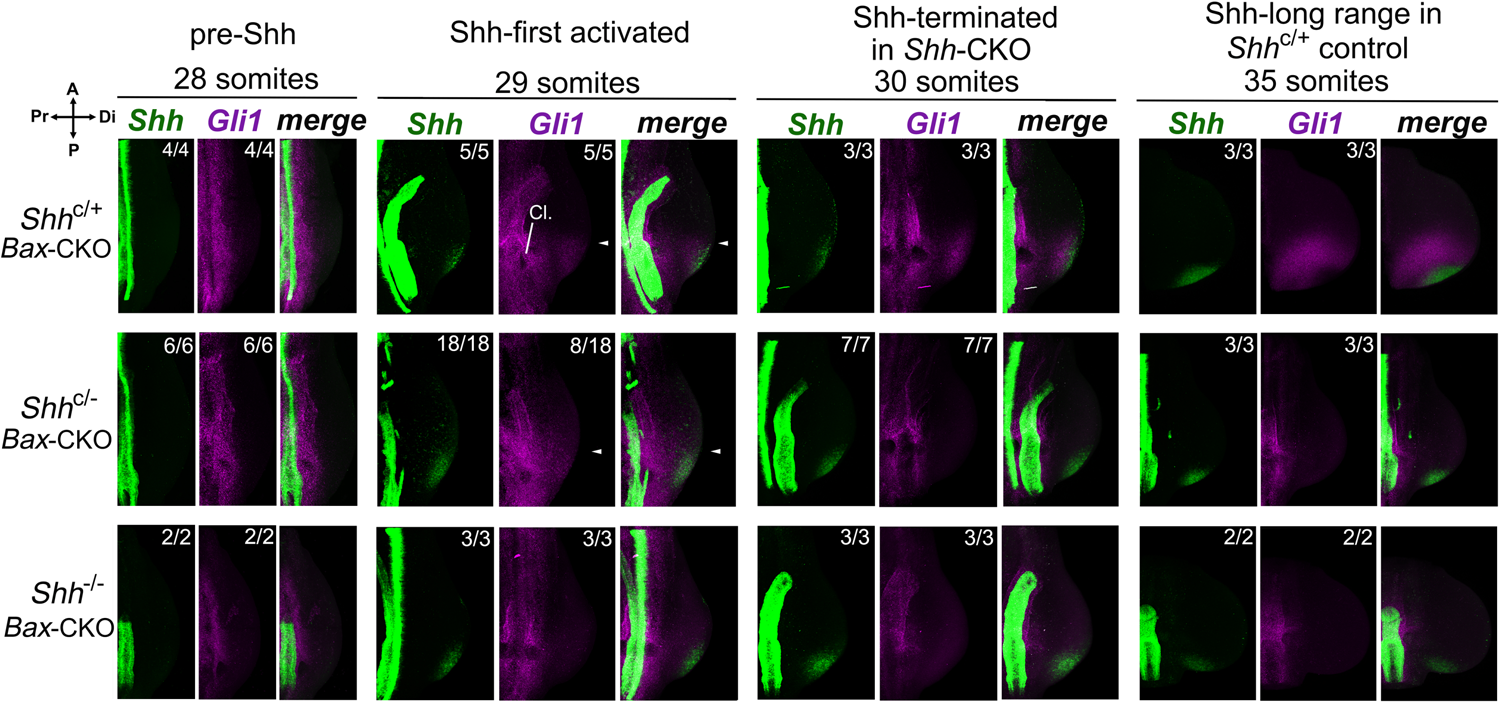
Only short-range Shh response is detected in *Shh*-CKO;*Bax*-CKO embryos during transient *Shh* expression window. *Shh* expression and activity was assayed by *Shh* (green) and *Gli1* (purple) RNA in situ HCR^16^ at somite stages indicated. After tamoxifen injection at E9.5+3h (as in Fig. 1B,C), Shh response (*Gli1*) was detected at the 29 somite stage, shortly after *Shh* expression onset, in all control *Shh*^+/-^;*Bax*-CKO (5/5) and in a subset of *Shh*-CKO;*Bax*-CKO (8/18) embryos. By 30 somites (2h later) and at 35 somites (about 12h later), Shh response became robust in all controls but was absent in all *Shh*-CKO embryos. A-P *Gli1* expression extent in distal limb bud is very similar to that of *Shh* in both the control and the *Shh*-CKO embryos during the transient expression window (29 somites; merged panels, arrows). Note that non-functional (exon 2-deleted) *Shh* RNA remains detectable in both late-stage *Shh*-CKO and in *Shh*^-/-^ null embryos (with absent *Gli1* RNA signal). Numbers analyzed with result shown are indicated at bottom of each panel, with remainder negative for expression. Cl, cloacal-urogenital region, which also responds to local Shh and Ihh signaling^18, 19^. Related to Figure 2.

### Digit Rescue in Shh-CKO by enforced cell survival is not a consequence of residual or re-activated Hh pathway function or pathway target de-repression

There are several alternative possibilities to account for the observed rescue of normal limb development in *Shh*-CKO;*Bax*-CKO embryos that need to be ruled out. First, a re-emergence of Hedgehog pathway activity, which might be the result of mosaic recombination of the *Shh*-floxed allele, de novo re-induction of a ZPA in a new cell population, or ectopic induction of an alternate (Ihh or Dhh) ligand, for example.

Each of these would result in, and be detected by, re-activation of direct Shh targets (*Ptch1, Gli1* RNA) that provide a more sensitive readout of pathway activation than measuring Shh-ligand expression. Monitoring of direct Shh target *Ptch1* (0/7, 0/9) and *Gli1* (0/7) RNAs at both early and late patterning stages, or inclusion of a *Gli1*^LacZ/+^ knock-in allele (0/8) to provide a highly sensitive Shh-response reporter with enzymatic amplification^20^ at early stages, both failed to detect any Hh pathway recovery in *Shh*-CKO;*Bax*-CKO embryos (Fig. 4A). In particular, mosaic recombination of the *Shh*-CKO allele would leave residual Shh-expressing cells that would proliferate and consequently be expected to increase the *Ptch* or *Gli1* reporter signal over time.

**Figure 4.**
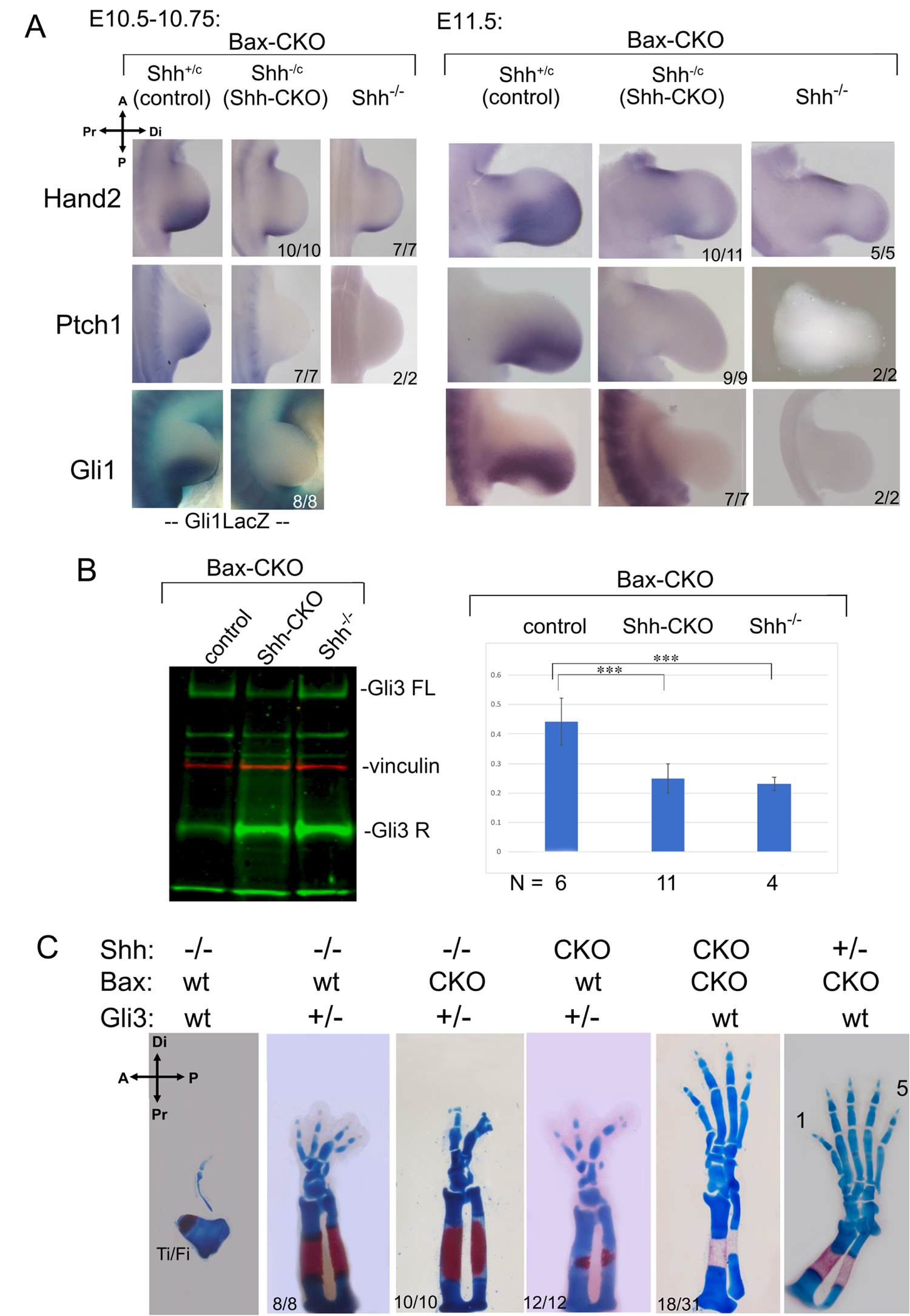
Digit rescue in *Shh*-CKO;*Bax*-CKO embryos is not due to Shh pathway re-activation. **(A)** RNA expression monitoring Shh pathway activity (*Hand2, Ptch1, Gli1*) and assay for LacZ activity from a *Gli1*^LacZ/+^ knock-in allele after *Shh* removal by tamoxifen at E9.5+3h (as in Fig. 1B). Mutant numbers analyzed with the result shown are indicated in each panel in **A**. In the remainder, expression is unchanged from the null *Shh*^-/-^;*Bax*-CKO. **(B)** Gli3 full-length (Gli3FL) and repressor (Gli3R) protein quantitation in E10.5 hindlimbs, following *Bax/Bak* and *Shh* removal as in Fig. 1B,C. Typical blot shown to the left. Gli3 FL/R ratios shown to right in bar-graph with N numbers analyzed for each genotype. Gli3 FL/R is equivalently reduced in *Shh*-CKO;*Bax*-CKO (p=0.00002) and in *Shh*^-/-^;*Bax*-CKO (p=0.001) compared to control, and there is no significant difference in Gli3 FL/R between *Shh*-CKO;*Bax*-CKO and *Shh*^-/-^;*Bax*-CKO (p=0.48). ***, p <0.001. **(C)** Effect of *Gli3* dosage reduction (*Gli3*^+/-^) on *Shh*-CKO and on *Shh*^-/-^ skeletal phenotypes (E16.5), compared to the effects of *Bax/Bak* removal (*Bax*-CKO).

Another major, important possibility to address, is potential loss of pathway repression by Gli3 repressor (Gli3R). Shh prevents processing of Gli2/Gli3 nuclear effectors from full-length activators (Gli2FL/Gli3FL; GliA) to truncated repressors (Gli2R/Gli3R) of Shh targets^21, 22^. Release from Gli3 repression arguably plays the main role in most Shh limb target regulation^13, 14, 23, 24^. Consequently, key limb target activation may occur ligand-independently, without activating target reporters (*Gli1, Ptch1*) via either Gli3R removal or functional antagonism. We used several approaches to test for evidence of altered net Gli3R activity. First, *Hand2*, which induces ZPA/Shh by antagonizing *Gli3*^23^, is directly repressed by Gli3R^15^ and remained absent from rescued *Shh*-CKO;*Bax*-CKO in both early (10/10) and later (10/11) stage limb buds (Fig. 4A). Secondly, Gli3R activity can be modulated at the protein level by altered processing or degradation^21, 22^. Although *Bax/Bak* removal in wildtype limb buds has no impact on skeletal patterning, an altered balance of Bcl2 family members has been reported to affect Gli-processing activity indirectly and could generate a Gli3R deficit^25^. We examined Gli3 protein levels in early individual limb buds (E10.75, Fig. 4B). Gli3FL/Gli3R ratios in *Shh*-CKO;*Bax*-CKO were unchanged from *Shh*^-/-^;*Bax*-CKO limb buds, and both were equally reduced compared to Shh-expressing controls (Fig. 4B), indicating that rescue was not explained by reduced Gli3R protein. Thirdly, “effective” repressor activity of Gli3R may be altered by interacting protein partners without changing quantitative protein level^26, 27^. To exclude altered net Gli3R activity by any mechanism, we used a genetic test to compare the phenotypic effect of rescuing cell survival in *Shh*-CKO;*Bax*-CKO with that of intentional *Gli3* dosage reduction. We compared *Bax/Bak* removal with *Gli3* dosage (*Gli3*^+/-^) effects in both *Shh*-CKO and *Shh*^-/-^ limbs. Complete *Gli3* loss alone rescues limb development in *Shh*^-/-^ embryos, albeit with synpolydactyly^13, 14^ and phalangeal morphology changes. However, haploid *Gli3* dosage (*Shh*^-/-^;*Gli3*^+/-^) has an intermediate effect on the *Shh*^-/-^ null phenotype^13^ (improved zeugopod morphology, several small digit rudiments; Fig. 4C, 8/8). In contrast to *Shh*^-/-^;*Gli3*^+/-^, no change in the *Shh*^-/-^ null phenotype was observed when *Bax/Bak* was removed (Fig. 1D; 18/18). Furthermore, removing *Bax/Bak* in the *Shh*^-/-^;*Gli3*^+/-^ limb did not improve limb skeletal phenotype beyond the effect of *Gli3* dosage reduction alone (Fig. 4C, 10/10), suggesting that *Bax/Bak* removal did not impact the “effective” net Gli3R level significantly and that reduced Gli3 dosage, even together with enforced cell survival, in the Shh null does not mimic, or substitute for, the effect of transient Shh expression with enforced cell survival (Shh-CKO;Bax-CKO).

Whereas *Bax/Bak* removal had little effect on the *Shh*^-/-^ null compared to *Gli3* dosage reduction, the reverse holds for the *Shh*-CKO. Without enforced cell survival (*Bax/Bak* removal), the *Shh*-CKO;*Gli3*^+/-^ was phenotypically identical to the *Shh*^-/-^;*Gli3*^+/-^ (12/12; Fig. 4C). In contrast, in *Shh*-CKO;*Bax*-CKO limbs with wildtype *Gli3* dosage (*Gli3*^+/+^), both zeugopod and between 3-5 digits with normal morphologies and clear A-P polarity were restored (18/31; Figs. 1D, 4C). Together, these results argue strongly against altered Gli3R activity *per se* as a mechanistic basis for the restoration of normally polarized limb development in the early *Shh*-CKO with enforced cell survival to bypass Shh late function.

### Expression of certain outgrowth and patterning regulators is sustained after Shh loss if the Shh requirement in cell survival is bypassed (Shh-CKO;Bax-CKO)

To assess whether downstream regulators of limb outgrowth and digit patterning are restored by transient Shh exposure, we examined expression of major downstream targets that regulate limb bud outgrowth (AER/Fgf signaling^28, 29^) and digit patterning (5’*Hox* genes^30, 31^). Unlike *Hand2* and direct GliA-regulated targets (*Ptch1*, *Gli1*), expression of key outgrowth and patterning regulators is maintained in a subset of *Shh*-CKO;*Bax*-CKO embryos, roughly correlating with the observed 50% occurrence of an early, transient Shh activity pulse, and subsequent skeletal rescue (Figs. 1, 5, S1-S3). In the remainder, expression profiles in *Shh-*CKO;*Bax-*CKO resembled the null *Shh*^-/-^;*Bax*-CKO (Figs. 5, S2, S3).

**Figure 5.**
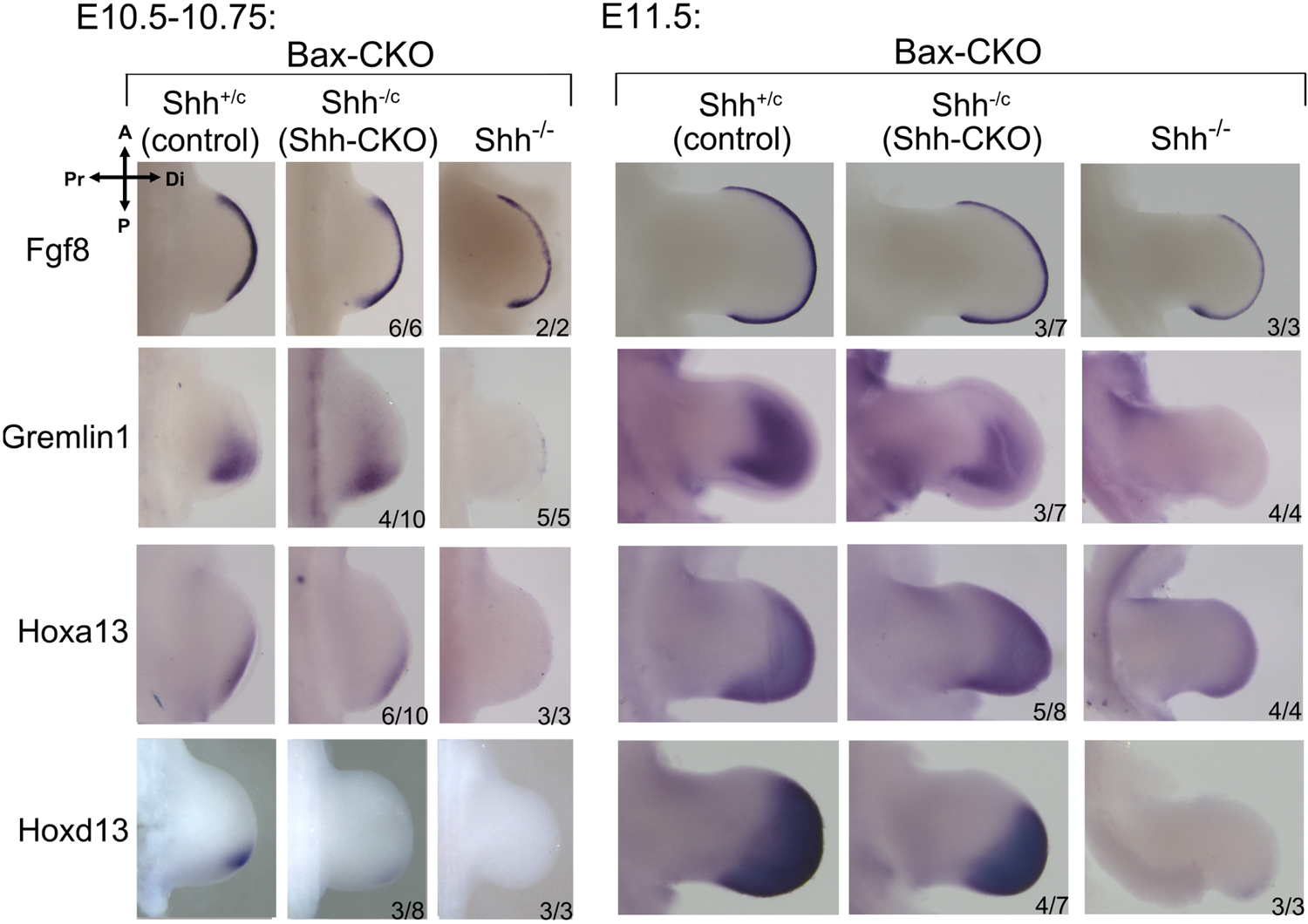
Expression of key targets implicated in outgrowth and patterning is maintained in *Shh*-CKO;*Bax*-CKO embryos. RNA expression monitoring Shh targets that regulate outgrowth (*Fgf8, Grem1*) and patterning (*Hoxa13, Hoxd13*) at early and later stages after *Shh* removal by tamoxifen at E9.5+3h (as in Fig. 1B and 2A). *Fgf8* expression is unaltered even in *Shh*^-/-^ null hindlimb at E10.75^2^, but was reduced in *Shh*^-/-^;Bax-CKO by E11.5. *Shh*-CKO;*Bax*-CKO embryo numbers analyzed with the result shown are indicated in each panel (preservation of expression). In the remainder, expression was unchanged from that observed in the null *Shh*^-/-^;*Bax*-CKO. See also Figure S2 and S3.

The direct Shh target *Grem1* plays a key role in AER/Fgf8 maintenance^29^ and consequently *Fgf8* expression declines in null *Shh*^-/-^ hindlimb after E10.5^2^. *Shh*^-/-^;*Bax*-CKO limb buds likewise lacked early and late *Grem1* expression (absent in 5/5, 4/4, Fig. 5) and by E11.5 *Fgf8* expression was clearly reduced (3/3, Fig. 5). In contrast, a subset of *Shh*-CKO;*Bax*-CKO embryos maintained early and late *Grem1* expression (4/10 and 3/7, Fig. 5), and preserved *Fgf8* after E10.5 (3/7, Fig. 5). The Fgf8-regulated target, *Cyp26b1*, required to clear retinoids for distal limb progression^32^, was also maintained in *Shh*-CKO;*Bax*-CKO (4/6) but declined in *Shh*^-/-^;*Bax*-CKO by E11.5 (3/3, Fig. S2).

Several 5’*Hox* genes regulate AP patterning downstream of Shh. *Hoxd13* and *Hoxa13*, critical for digit specification^31^, are both expressed only at very late stages and at low levels in *Shh*^-/-^;*Bax*-CKO (3/3, 4/4) compared to controls (Fig. 5). Low level distal *Hoxa13* expression was already detected early in a subset of *Shh*-CKO,*Bax*-CKO hindlimb buds (6/10), and became robust at later stages (5/8, Fig. 5). *Hoxd13* was detected at a trace level early (3/8), but was clearly detectable at the onset of the second phase 5’*Hoxd* distal footplate expansion^33^ (4/7, E11.5, Fig. 5), and well prior to the late condensation stage after all digit rays have formed when both the *Shh*^-/-^ null^2^ and the *Shh*^-/-^;*Bax*-CKO re-express *Hoxd13* (∼E12.5, Fig. S2). Since 5’*Hoxd* genes act mainly during the distal expansion phase to determine digit identity by regulating late interdigit signaling centers^34^, this sustained second phase *Hoxd13* activation likely suffices for the morphogenesis of normal distinct digit types in *Shh*-CKO;*Bax*-CKO compared to *Shh*^-/-^;*Bax*-CKO (null) embryos. *Hox11* paralogs play a key role in zeugopod patterning and growth^30^, which is also highly perturbed in the *Shh*^-/-^ but restored in *Shh*-CKO;*Bax*-CKO hindlimbs (tibia/fibula, Fig. 1D, 4C). Early *Hoxd11* expression was similar to controls even in *Shh*^-/-^;*Bax*-CKO (2/2), but became undetectable by the second phase distal expansion (2/2). In contrast, *Hoxd11* was maintained at control levels both early and late in a subset of *Shh*-CKO;*Bax*-CKO limb buds (5/5, 2/7, Fig. S2), consistent with zeugopod rescue frequency. Shh inhibition in short term mouse limb bud cultures^24, 35^ also suggests that some direct Shh targets are maintained if Shh activity is curtailed after onset, as we have shown here (*Grem1*; *Jag1* in Fig. S2), but in those studies other downstream targets, particularly 5’*Hoxd* genes, appeared to require sustained Shh activity for their continued expression. However, development does not progress normally in short-term ex vivo cultures, precluding analysis of the second phase 5’*Hoxd* expression (which is selectively restored in the *Shh*-CKO;*Bax*-CKO) following early Shh inhibition in cultured limb buds.

We also examined the expression of anterior regulators that are repressed/antagonized by Shh activity in the normal posterior limb bud. Unlike the posterior regulators, major anteriorly expressed regulators of patterning, *Irx3*^36, 37^ and *Alx4*^14^, are only modestly altered in their early extent even in the *Shh* null (*Shh*^-/-^) mutant (Fig S3). However, the slight posterior expression extension present in *Shh*^-/-^ was also observed in about 50% of the *Shh*-CKO;*Bax-*CKO embryos (Irx3 2/4; Alx4 5/8). Our results indicate that expression of targets important for both limb bud outgrowth and patterning are sustained by a transient pulse of Shh with enforced cell survival, providing a basis for the phenotypic rescue of limb development, but do not address the issue of how transient Shh activity leads to stable, AP-graded expression of certain Shh regulated targets in the limb.

### Shh is required indirectly to specify digit 1 (thumb)

To test if Shh acts via a relay mechanism, we used a genetic strategy (Fig. 6A) to activate Shh targets autonomously only in ZPA cells and ask if any non-ZPA-derived digits are rescued in the complete absence of Shh ligand (*Shh* null). Cell-autonomous pathway activation was achieved using a conditional transgene (*Rosa*^SmoM2^), expressing a constitutively-active form of Smoothened (SmoM2)^38^, a membrane GPCR essential for transducing Shh^39^. The *Shh*^Cre^ knock-in allele^4^ was used to restrict SmoM2 and Shh target activation to ZPA cells (*Shh*^cre^;*Rosa*^SmoM2/+^; referred to as Shh-SmoM2+), and evaluated in the *Shh* null background (*Shh*^cre/-^). Enforced Shh-response by SmoM2 in the ZPA also affects *Ihh* targets and chondrogenesis^40^, precluding morphologic evaluation of any ZPA-descended digits (d4,d5 in wildtype), but differentiation of other digits is unaffected (see Fig. S4A). Unexpectedly, in both *Shh*^cre/-^ fore- and hindlimbs, enforced cell-autonomous pathway activation of *Shh*^cre/-^;Shh-SmoM2 in the ZPA rescued formation of a morphologically normal biphalangeal digit 1 at high frequency (66%; 21/32 limbs, Fig. 6B). Digit 1 specification is thought to be Hh-independent, based on a persistent d1-like structure in *Shh* null hindlimbs^2^ and the normal lack of direct Shh-response in the d1 progenitor territory^9^ (see also Fig. 2). In fact, direct Shh-response has been shown to inhibit d1 territory formation^36^. The rescued d1 in *Shh*^cre/-^;Shh-SmoM2 was not ZPA-derived (LacZ reporter-negative), but arose entirely from anterior limb bud progenitors (5/5; Fig. 6C) at the anterior limb margin where d1 normally forms (see Fig S4F), and was not the result of either cryptic Hh ligand or downstream pathway activation as seen in a small fraction of control Shh-SmoM2+ limb buds (Gli1, 0/8; Ptch1, 0/10; Fig. S4A,C). In contrast, we found that the single dysmorphic digit which forms in the *Shh* null mutant (*Shh*^cre/-^) is actually entirely descended from posterior ZPA (LacZ+) d4/d5 progenitor cells (Figs. 6C, S4A), which is consistent with the extensive/broad anterior apoptosis present in *Shh*^-/-^ limb buds. Unlike the ZPA-origin of the *Shh* null digit, our lineage analysis clearly demonstrates that the morphological d1 rescued in the *Shh*^cre/-^;Shh-SmoM2 arises from anterior non-ZPA limb cells. *Hoxa13*, which is uniquely essential for d1 specification^31, 41^, and is absent or greatly reduced in *Shh*^Cre/-^ null (0/4, Fig. 6D, forelimb shown), was restored across the *Shh*^cre/-^;Shh-SmoM2+ distal limb bud (6/6; Fig. 6D), confirming d1 identity. Likewise, a late d1-specific marker (*Uncx4.1*) was also expressed in the rescued digit domain of *Shh*^cre/-^;Shh-SmoM2 (6/8), but not in *Shh*^Cre/-^ (0/4; Fig. S4B). Together, these results strongly argue that a bona fide digit 1 is absent in the *Shh* null mutant and is restored in the *Shh*^cre/-^;Shh-SmoM2, indicating that d1 specification is actually Shh-dependent and requires an indirect signal activated downstream of Shh-response in the ZPA.

**Figure 6.**
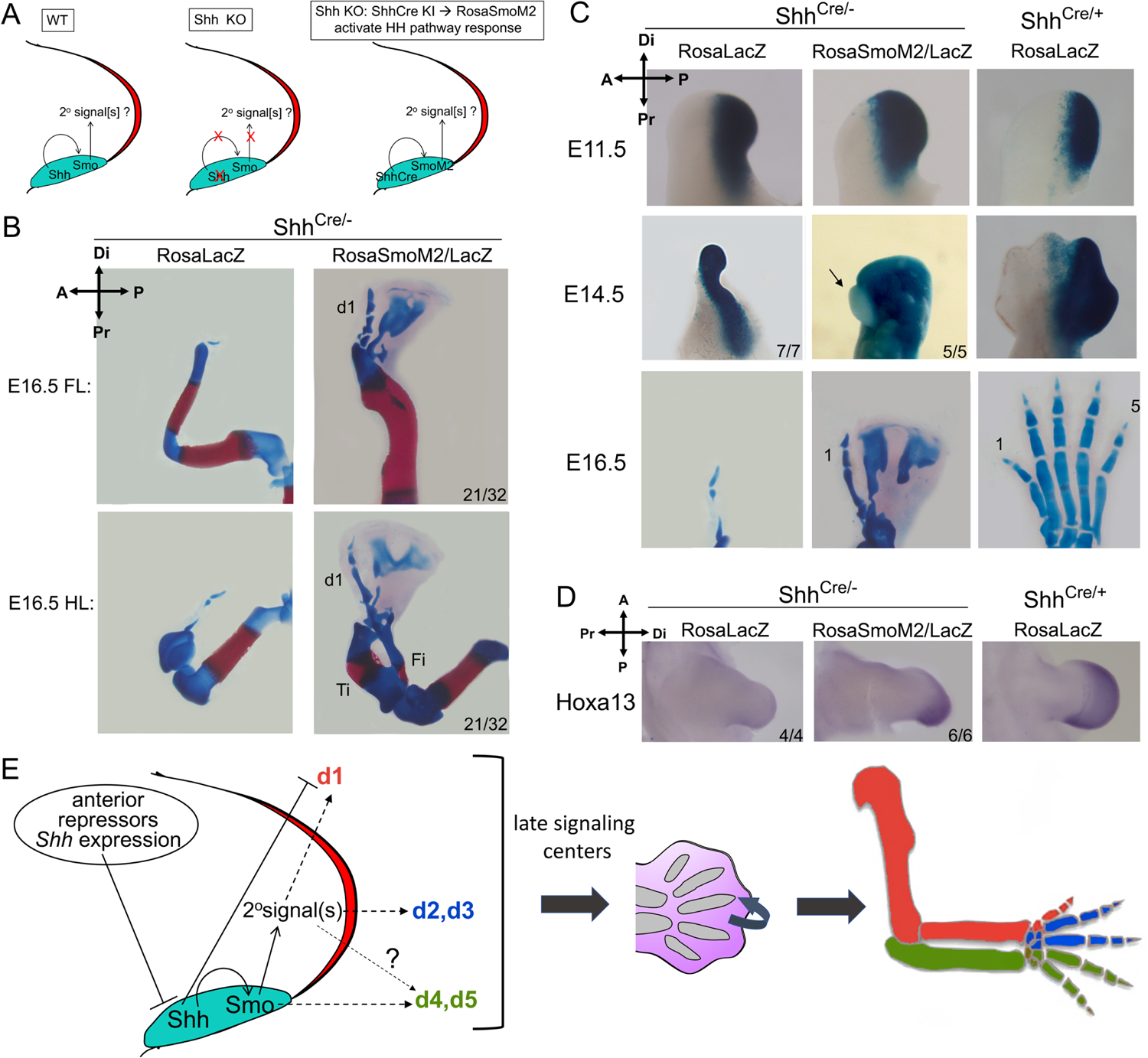
Relay signals downstream of enforced Shh-response in ZPA restore normal digit 1 in *Shh*^-/-^ null limb. **(A)** Activation of Shh-response in ZPA of null limb bud (*Shh*^Cre/-^) by SmoM2 (*Shh*^cre/-^;Shh-SmoM2+). Any effect on non-ZPA digits requires non-autonomous relay signal(s). **(B)** Skeletal stain showing normal digit 1 (d1) in *Shh*^cre/-^;Shh-SmoM2+ forelimbs and hindlimbs (21/32). Ti, tibia; Fi, fibula. **(C)** ZPA-lineage analysis and close-up of mutant footplate with activated SmoM2. The anterior-most digit in *Shh*^cre/-^;Shh-SmoM2+ remains devoid of LacZ+ cells (5/5, arrow), whereas *Shh* null digit arises entirely from LacZ+ ZPA-descended cells (7/7). **(D)** *Hoxa13* RNA in E11.5 forelimb is restored in *Shh*^cre/-^;Shh-SmoM2+ (6/6), but absent in *Shh* null (4/4). **(E).** Model for digit specification by Shh via an indirect relay mechanism (see text discussion). Transient Shh specifies non-ZPA digits (d1-d3) indirectly by activating relay signals in ZPA. Direct Shh signaling selectively inhibits^36^ but indirect relay signals promote d1 specification, establishing a unique d1 (thumb) regulatory hierarchy. Relay signaling, initiated by early transient Shh, ultimately sets up late interdigit signaling centers to regulate final digit identity. See also Figure S4.

Why was only d1, but not other non-ZPA digits (d2,3) restored by Shh-SmoM2? Shh cell survival function remains absent and *Shh*^cre/-^;Shh-SmoM2+ limb buds display considerable anterior apoptosis (Fig. S4D-F), but enforcing cell survival (with *Bax*^-/-^;*Bak*^-/-^, see Table 1) did not rescue any further digit formation besides d1 (8/8). We suspect that the failure to rescue d2-3 reflects a problem inherent in the timing of enforced SmoM2/Shh-response in ZPA cells (which is induced by *Shh*^Cre^ only <u>after</u> normal *Shh* onset). The delayed specification of d1 relative to other digits^1, 41^ would be consistent with this observed, selective d1 rescue.

## Discussion

Our genetic rescue reveals two distinct roles for Shh in the limb: a transient (2-3hr), early requirement that is critical to specify all digits, and a sustained requirement to promote cell survival. This transient Shh pulse is both necessary and sufficient for normal limb morphogenesis, if the role of Shh in maintaining cell survival is bypassed (by *Bax/Bak* removal). Yet genetic lineage tracing shows that Shh acts only short-range and Shh-response is restricted to ZPA-derived digit progenitors (d4-d5) during the same time interval that suffices to specify all digits, and HCR confirms that the transient signal response in the *Shh*-CKO mutant is short range and restricted to the same limited zone as seen in the wildtype limb bud during the same time interval. Both biochemical and genetic assays reveal that the Gli3R activity gradient is unperturbed by the short Shh activity pulse, as expected in the absence of any long-range Hh response. Together, these results indicate that Shh is not a limb morphogen but acts via a signal relay mechanism to specify at least the non-ZPA digits (d1-3, Fig. 6E). Our results suggest that Shh may act locally to specify d4/5 directly, but don’t exclude an indirect contribution. Digit 1 specification, occurring relatively late, is further distinguished by being repressed by direct Shh response, yet Shh-dependent indirectly (discussed below). These different responses to Shh signaling define up to three distinct types of regulation that lead to the specification of posterior ZPA-descended digits (4-5), anterior digits (2-3) and digit 1 (thumb; see Fig 6E). Unexpectedly, long-range Shh signaling which occurs only at later stages, when the limb bud is expanding, plays a critical role in sustaining cell survival across the d2-5 territories, but not in patterning directly.

We carefully considered, and tested for, alternative explanations for these results that would invalidate our conclusions, either because of persistent pathway activity over a more extended time frame that escaped detection, or because of a long-range response during the early *Shh* expression pulse that escaped detection.

A low-level persistence of Shh pathway activation, owing to either incomplete recombination to delete Shh during the period of tamoxifen activity or to some compensatory feedback leading to ectopic Shh activation or activation of another Hh ligand should all result in the continued activation of direct targets *Gli1* and *Ptch1*, which are detectable at much lower levels than is *Shh* RNA^5, 12, 42^. Furthermore, because Shh-expressing cells proliferate in the early limb bud (later giving rise to posterior digits), the activation of direct target reporters should become increasingly apparent over time (E10.5-11.5). Yet several methods to visualize response at later stages, including HCR with RNA hybrid amplification or a Gli1LacZ reporter with enzymatic amplification, did not detect the presence of Shh direct-target response in any *Shh*-CKO;*Bax-*CKO limbs. Target RNAs became undetectable within 3 hrs of *Shh* expression onset (28/28 by ∼E10+3, 30 somites; Figs. 1C, 3, S1A), despite increasingly strong expression in all control (*Shh*+) embryos, and remained completely negative at 12 hrs later (26/26 total at E10.5/10.75; Figs. 1C, 3, 4A, S1A), and at 36 hrs (16/16 total at E11.5; Fig. 4A), times by which target *Gli1* and *Ptch1* RNA expression has become broad and abundant. In addition, the phenotype of *Bax*+ *Shh*-CKO sibling embryos (see Fig. 1) was uniformly identical to the *Shh* germline null (28/28 embryos total, Fig. 1D, Table 1), which is incompatible with trace level residual Shh signaling. It has been shown that persistent, very low level Shh signaling (generated by deleting *Shh* with *ShhCre*, because of a positive feedback loop inducing ZPA cells) results in a very mild digit phenotype (16/16) despite only trace level *Shh* expression and modest *Ptch1* response^7^.

Failure to detect long-range signaling response triggered during the time window of transient Shh activity in the *Shh*-CKO;*Bax*-CKO could result from either a difference in the response-range between the mutant and controls such that signaling response in the mutant extends farther than the lineage tracing of response in wildtype controls would suggest, or from undetected effects of long-range signaling on Gli3R activity level and target de-repression. Lineage tracing analysis on Shh-responding cells in control (wildtype) limb buds clearly showed no response beyond the ZPA region (future digit 4-5 territory) during the equivalent time frame of transient Shh activity in the rescued *Shh*-CKO;*Bax*-CKO and in fact long-range response was not detected until E10.25 and clearly evident at E10.5, despite sensitive enzymatic amplification (LacZ+ reporter recombinants). These tamoxifen treatment times for long-range response detection are considerably later than that used to rescue digit formation in the *Shh*-CKO. Such lineage analysis cannot be examined directly in the *Shh*-CKO;*Bax*-CKO without developing a suite of non-Cre/loxp based conditional alleles and deletion drivers. Therefore, we evaluated the extent of Shh response in *Shh*-CKO;*Bax*-CKO limb buds in comparison to wildtype control more directly, using HCR to simultaneously visualize both *Shh* and *Gli1* in the same limb bud during the transient Shh activity period. These results reveal Shh-response only in cells within and narrowly surrounding the ZPA region, with an identical spatial A-P extent in both controls and in *Shh*-CKO;*Bax*-CKO, confirming that only short-range Shh signaling is present during the transient Shh activity phase in the rescued *Shh*-CKO;*Bax*-CKO.

Long-range effects of Shh on Gli3 repressor levels (Gli3R) and target gene “de-repression” are not directly measured by GliA-dependent target reporters (*Gli1, Ptch1*), although many Shh targets are regulated mainly by de-repression^24, 43^. However, there is no single, uniform expression response for this target class, because their regulation by other activating transcription factors varies highly from gene to gene^24^, and consequently there are no good universal “reporters” for response to Shh-induced de-repression. We used a genetic test modulating Gli3 dosage to assess whether the *Shh*-CKO;*Bax*-CKO rescue results might be caused by altered Gli3R activity level. Phenotypic effects of Gli3 dosage modulation also serve as a gauge to assess potential change in the anteriorly-biased Gli3R gradient caused by any long-range signaling response and target de-repression during transient Shh activity. Our results (Fig. 4C) indicate that *Gli3* dosage reduction (*Gli3*^+/-^) does not phenocopy the digit rescue by enforced cell survival in the *Shh*-CKO, and has no impact on the *Shh*-CKO skeletal phenotype beyond the *Gli3*^+/-^ effect in the *Shh*^-/-^ null mutant, arguing against long-range Shh-signaling modulation of Gli3R activity in the rescued *Shh*-CKO.

Digit identity is morphologic in nature, arising via distinct organizations of the same tissues and not based in cell fate changes, features suggesting progressive specification. Indeed, work in both chick and mouse indicates that late interdigit signaling centers^34, 44–47^ impinge on digit tip progenitors to regulate phalanx numbers formed, determining final digit identities. Shh may regulate digits specified at particular A-P limb positions via relay signals to establish such late signaling centers (Fig. 6E). A relay mechanism is incompatible with Shh acting either as a traditional morphogen or by temporal integration, as concluded from some chick work^3, 6, 7^. These studies rely on pharmacologic inhibition that may persist to later stages to affect Ihh in digit tip progenitors, leading to phalanx loss^48^ that may score as digit identity changes attributed to Shh inhibition; reduced cell survival by Shh suppression may also impact late-stage digit morphogenesis. Notably, other chick studies have suggested involvement of downstream relay signals^6, 49, 50^, but did not discriminate if Shh also acts as a morphogen. Whether apparent mouse-chick differences in Shh function reflect biologic or technical factors remains to be explored.

Uncovering the mechanisms that sustain stable gene activation after only transient Shh exposure will be an important future focus to illuminate how Shh patterns the developing limb. Stable alterations in target gene expression following transient Shh exposure could involve several, non-mutually exclusive mechanisms including chromatin and/or DNA modifications and relay mechanisms incorporating lock-on circuitry^51^. Recent work suggests that Shh targets are poised for expression as soon as Gli3-mediated repression is alleviated^43^, and a transient burst of Shh activity could trigger a self-reinforcing bi-stable switch^51^ if activating factors that reinforce target expression also block re-introduction of repressive marks by Gli3R. Furthermore, our results clearly implicate non-cell autonomous relay signals that act downstream of transient Shh activity to specify non-ZPA derived digits. Such relay signals can become rapidly self-sustaining via feedback loops, as occurs with Fgf10-Fgf8 signaling downstream of transient Tbx5 activity in the limb^52, 53^.

An alternative to a relay mechanism for patterning digit territories not directly responsive to transient Shh signaling would be the maintenance of polarity, once initiated by transient Shh activity, at the level of antagonistic interactions between “posterior” (SHH target) and “anterior” transcription factors, which are initially co-expressed in very early-stage limb bud mesenchymal cells^54^. However, such a scenario is difficult to reconcile with the finding that A-P polarized expression of some, but not other, targets is maintained in the rescued *Shh*-CKO;*Bax*-CKO limb buds and is accomplished in the absence of any *Hand2* expression, which normally antagonizes anterior Gli3R to maintain posterior limb bud asymmetry both upstream and downstream of Shh^23^. Furthermore, lineage tracing indicates that the response to “transient” Shh is entirely restricted to the posterior ZPA region, which would generate clear-cut anterior and posterior limb bud compartments, rather than the graded A-P target gene expression seen for some target genes maintained in the rescued *Shh*-CKO;*Bax*-CKO (eg. *Grem1, Hoxa13*, Fig. 5). Spatially graded target expression would require some type of non-autonomous effect, whether induced by Shh effectors (Gli2/3) directly or by target transcription factors in the ZPA acting downstream of short-range Shh response.

Our genetic relay signal assay, in which Shh response was selectively enforced only in the ZPA region of the Shh null (*Shh*^-/-^) mutant, unexpectedly revealed that d1, like other anterior digits (d2-3), is also Shh-dependent. The rescued digit had the features of bona fide digit 1 (d1) based on both its positional and on expression/morphologic criteria. Lineage analysis showed origin entirely from non-ZPA cells (Fig. 6C), indicating d1-d3 progenitor origin, and restoration of cell survival selectively only in the very anterior-proximal mutant limb bud border is most consistent with d1 position (Fig. S4F). The rescued digit always has clear biphalangeal morphology and late marker expression associated with d1. Notably, distal-anterior *Hoxa13* expression which is uniquely required for digit 1 specification^41^, is also restored. Yet previous work has shown that direct Shh signaling selectively prevents formation of d1 territory, and indeed, a complex regulatory circuit repressing *Shh* expression-response anteriorly is required to specify d1^1, 36^. Unlike other digits specified by Shh-induced relay signals, the d1 territory remains outside the range of Shh signaling-response (eg. Fig 2B) during the entire expansion phase when Shh maintains cell survival and consequently d1 survival must be sustained differently. Why impose such a complex regulatory hierarchy for d1 specification (see Fig. 6E), involving repression by direct Shh signaling but requiring an indirect Shh-dependent relay signal? We propose that the unique control and consequent delay in d1 specification and growth/expansion enabled its independent evolution by uncoupling its regulation and morphogenesis from that governing other digits. Such regulatory uncoupling would facilitate the evolution of an opposable thumb, as well as other grasping/clutching adaptations important for arboreal tetrapods, including birds and mammals.

## Acknowledgements

We thank Cliff Tabin and Marian Ros for stimulating discussions and critical reading of the manuscript; Heinz Arnheiter, Denis Duboule, Bernhard Herrmann, Alex Joyner, Marie Kmita, Peter Koopman, Gail Martin, Marian Ros, Scott Stadler and Yingzi Yang for providing probes; and Sohyun Ahn, Chin Chiang, Alex Joyner, Andy McMahon, Cliff Tabin, Heiner Westphal and Yingzi Yang for providing mouse lines.

## Author Contributions

SM and JZ designed the project and wrote the paper, and JZ, RP, AT, and BDH performed the experiments.

## Data and materials availability

All data is available in manuscript and supplementary materials.

## Materials and Methods

### Mouse lines and tamoxifen injection

All animal studies were carried out according to the ethical guidelines of the Institutional Animal Care and Use Committee (IACUC) at NCI-Frederick under protocol #ASP-17-405. The *Shh*-floxed^12^, *Shh* null^55^, *Bax*-flox;*Bak*^-/-^^56^, *Gli1*^LacZ/+^^20^, and *Gli3* (*Xt-J*)^57^ mutant lines, and the *Hoxb6*CreER^11^, *Shh*Cre^4^, *Gli1*CreER^9^, *Rosa*SmoM2^38^, *Rosa*LacZ^58^, and *Rosa*EYFP^59^ mouse lines were all described previously. *Bax*-deleted (*Bax*^+/-^) mice were generated by crossing *Bax*-flox males with *Prrx1*Cre^60^ females to produce germ-line recombination. A detailed summary of the crosses used to generate embryos for different experiments and outcomes is provided in Table 1. For timed matings, noon on the date of the vaginal plug was defined as E0.5. For phenotypic rescue with *Hoxb6*CreER, pregnant mice were injected intraperitoneally with a single dose of 3mg tamoxifen (Sigma, T-5648) and 1mg progesterone^61^ (Watson, NDC 0591-3128-79) at E9.5+3hrs and embryos were collected at times indicated. For lineage tracing with *Gli1*CreER, a single dose of 0.5-1mg tamoxifen was injected at the times indicated.

### Whole mount in situ hybridization

Hybridizations were carried out following a previously described detailed protocol^62^. Embryos were fixed in 4% paraformaldehyde overnight, washed in PBS, gradually changed to absolute methanol and bleached with 5% H_2_O_2_. Embryos with different genotypes were treated together, in one tube, with 20ug/ml proteinase K for 8-16 mins based on the embryo age (this step was omitted for *Fgf8* probe). Gene-specific, digoxigenin-UTP labeled probes were synthesized from cDNA templates, and incubated with embryos in hybridization buffer with 50% formamide overnight at 70°C. The embryos were then washed with a series of buffers and incubated in alkaline-phosphatase-conjugated anti-digoxigenin antibody overnight at 4°C. After washing with 0.1% Tween in Tris buffered saline, embryos were incubated in BM purple (Sigma, 11442074001) or in NBT/BCIP to detect hybridized RNA.

### Hybridization chain reaction (HCR) *in situ* analysis

Whole mouse embryos at ages indicated were each fixed in 4% w/v paraformaldehyde in PBS overnight at 4°C, washed, bleached in 5:1 methanol/30% w/v hydrogen peroxide for 15 min. at 25°C and stored in methanol at −20°C until hybridization. Whole mount *in situ* hybridization of limb buds at ages indicated was performed using recommended conditions and solutions for third generation hybridization chain reaction (HCR) probes^16^ designed by Molecular Instruments (Los Angeles, CA) and analyzed by confocal microscopy. Maximum projection images of fluorescence intensity from confocal image stacks spanning the entire dorsoventral limb bud thickness were generated to compare the A-P extent of Shh and Gli1 RNA signals.

### Skeletal staining

For skeletal staining^63^, embryos were collected at E15.5-E18.5 and fixed in absolute ethanol overnight, followed by acetone overnight, and by staining in alcian blue/alizarin red in 95% ethanol overnight. After clearing in 1% KOH in H_2_O for several hours followed by 1% KOH in 20% glycerol, embryos were stored and imaged in 50% glycerol.

### Lysotracker staining

Embryos were collected in PBS and immediately incubated in lysotracker red (Sigma) in PBS with calcium and magnesium for 30 mins at 37°C. Embryos were then washed in PBS and fixed in 4% paraformaldehyde overnight at 4°C. Embryos were washed in PBS, transferred to absolute methanol in graded steps and cleared in benzyl alcohol/benzyl benzoate (BABB) solution to visualize staining.

### Western blot and quantification analysis

For western blot analysis, hindlimb buds from individual E10.75 embryos were dissected and lysed in 1x NuPAGE LDS sample buffer with 2% SDS and proteinase inhibitors. Reducing agent was added and samples were heated to 95 °C for 10 mins before loading. Two hindlimb buds (from 1 embryo) were loaded per lane, and electrophoresed in NuPAGE 3-8% Tris-Acetate protein gels. Proteins transferred to nitrocellulose membranes were probed with either affinity-purified polyclonal rabbit anti-Gli3^26^ or goat polyclonal anti-Gli3 (R&D, AF3690) and mouse anti-vinculin (Sigma, V9264) and visualized with fluorescent secondary antibodies (LI-COR IRDye 800CW, 926-32211, anti-rabbit green; 926-32214, anti-goat green; and with #680RD, #926-68072, anti-mouse red) using LI-COR Odyssey CLx. Bands were quantified with Image Studio software. For statistical analysis of western data (Gli3 FL/R), standard 2-sided t-test was used to calculate p values.

### Beta-galactosidase (LacZ) staining

Embryos were fixed in 2% paraformaldehyde with 0.2% glutaraldehyde for 1h, washed in PBS with 0.1% Tween and stained with XGal (1mg/ml) in PBT and 2mM MgCl_2_, 5mM Ferro-CN, 5mM Ferri-CN, at 37 °C for several hours.

## Supplemental Figure legends

**Figure S1.**
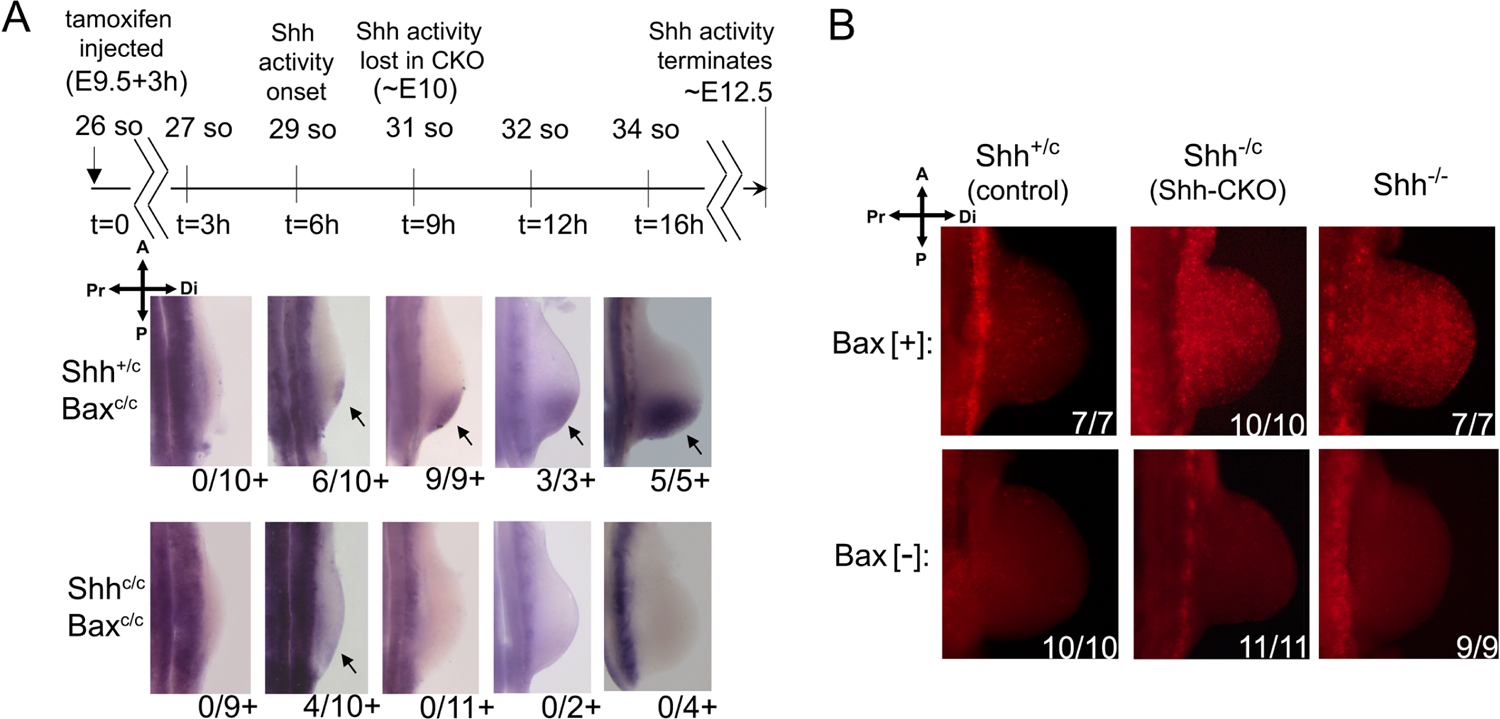
(related to Figure 1). Duration of Shh activity and enforced cell survival in *Shh*-CKO;*Bax*-CKO embryos. **(A)** Assay of Shh activity (direct response) by *Gli1* RNA expression after tamoxifen treatment (at times post-treatment indicated by timeline) in control (*Shh*^+/c^;*Bax*-CKO, upper panels) and in *Shh*-CKO;*Bax*-CKO hindlimb buds (lower panels). Note that Shh expression initiates at about 6-8 hrs <u>after</u> the time of tamoxifen injection. No activity is detected in either control (n=10) or *Shh*-CKO;*Bax*-CKO (n=9) hindlimbs at 3h after tamoxifen dosage. Shh activity was first detected at 6h after tamoxifen injection in a subset of control (6/10, arrow) and *Shh*-CKO;*Bax*-CKO (4/10, arrow) hindlimb buds, and became consistent and strong by 9h (n=9) and later (arrows) in control, but was absent in all *Shh*-CKO;*Bax*-CKO embryos (n=11) at 9h and later. Limb buds in all panels oriented with anterior at top and distal at right. so, somite. **(B)** Lysotracker staining for cell death at E10.75, following *Bax/Bak* and *Shh* removal with tamoxifen at E9.5+3h (as in panel **a**). In control hindlimbs with *Bax/Bak* function present (Bax [+], *Bax*^+/C^; upper panels) all *Shh*-CKO embryos (10/10) have extensive apoptosis at the same level as *Shh* null mutant (*Shh*^-/-^, n=7). In hindlimbs with *Bax/Bak* function deleted (Bax [-], *Bax*^C/C^; lower panels), no apoptosis is detected in either *Shh*-CKO (11/11) or in *Shh*^-/-^ null (9/9) hindlimb buds. Note that *Hoxb6CreER* is not expressed in somites^11^, where apoptosis remains present. Compass indicates limb bud orientation in all panels.

**Figure S2.**
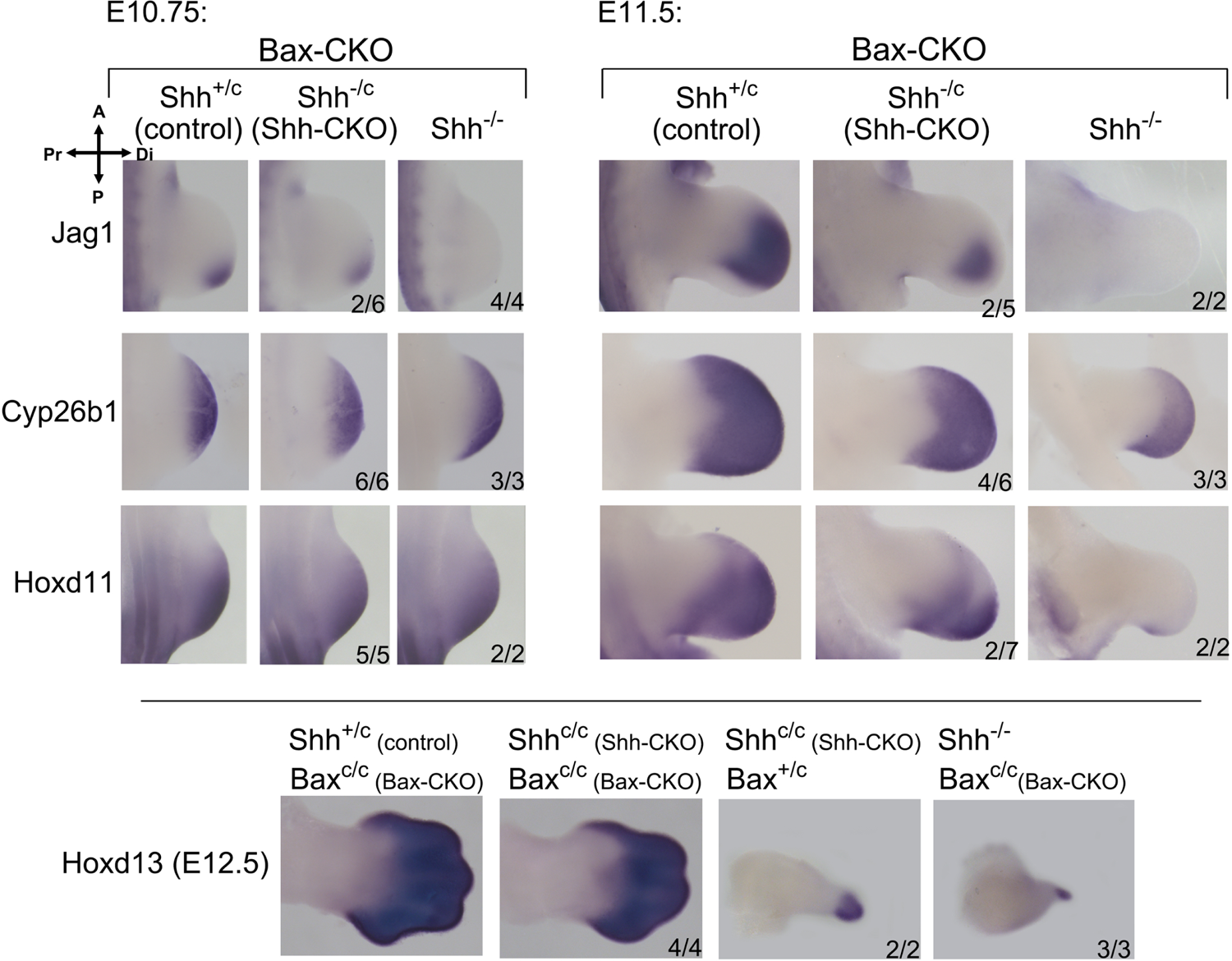
(related to Figure 5). Expression of Shh target genes implicated in limb bud outgrowth and digit patterning is maintained in *Shh*-CKO;*Bax*-CKO. *Jag1*, *Cyp26b1*, and *Hoxd11* expression at E10.75 and E11.5 are sustained in a subset (about 50%) of *Shh*-CKO;*Bax*-CKO hindlimbs. In null *Shh*^-/-^;*Bax*-CKO limbs, *Cyp26b1* and *Hoxd11* expression were preserved at E10.75, but markedly reduced or lost by E11.5. Mutant numbers analyzed with the result shown are indicated in each panel. In remaining *Shh*-CKO;*Bax*-CKO embryos, expression was unchanged from *Shh*^-/-^;*Bax*-CKO. The lower set of panels show *Hoxd13* expression at E12.5, which is maintained in *Shh*-CKO;*Bax*-CKO with phenotypic rescue of footplate (4/4), and in the digit rudiment of litter mate *Shh*-CKO;*Bax*^+/-^ hindlimbs (2/2) similarly to the null *Shh*^-/-^;*Bax*-CKO. Compass indicates limb bud orientation in all panels.

**Figure S3.**
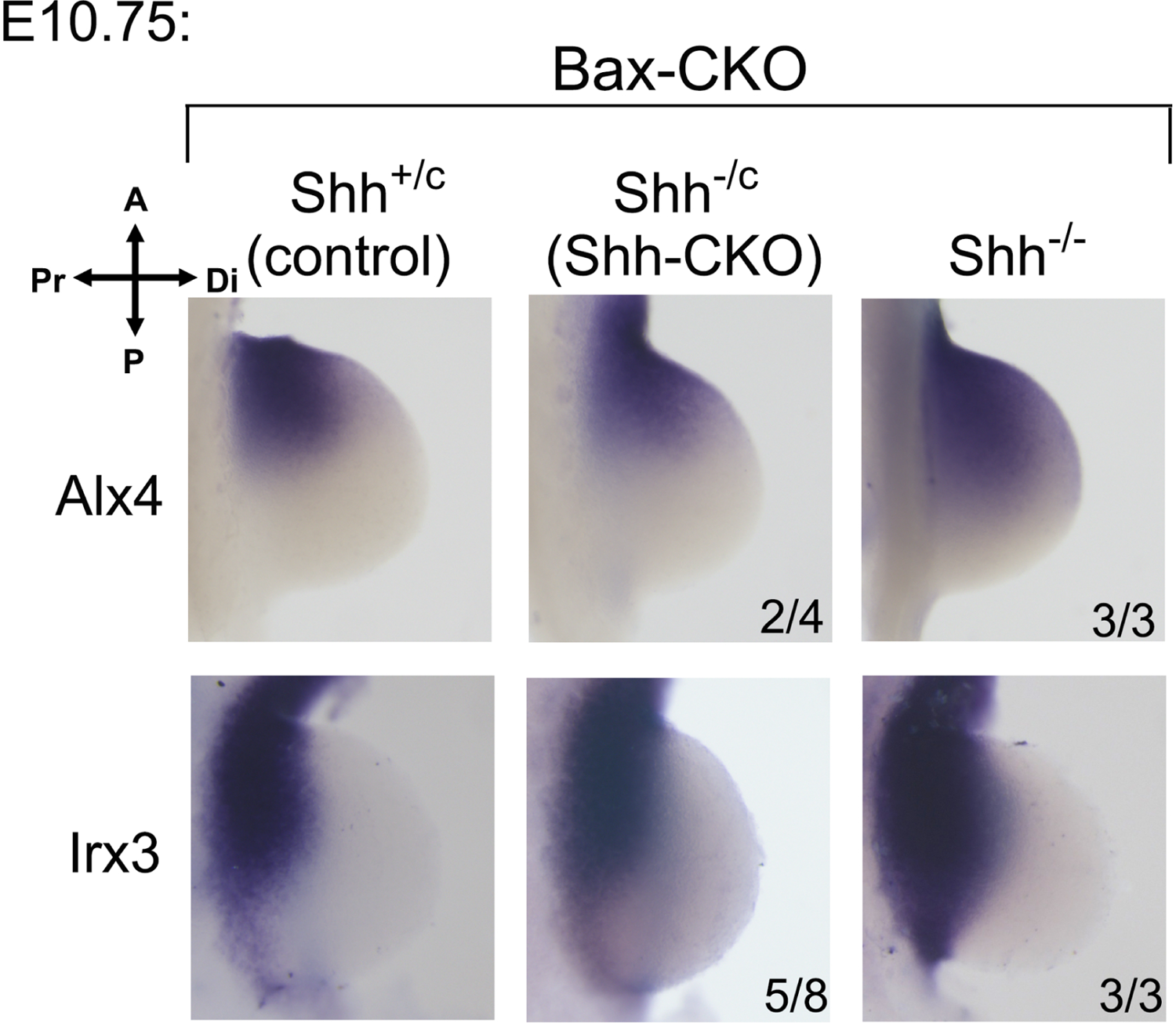
(related to Figure 5). Expression of anterior limb bud patterning regulators Alx4 and Irx3 is maintained in *Shh*-CKO;*Bax*-CKO. *Alx4* and *Irx3* expression at E10.75 remain anteriorly restricted similar to control limb buds in a subset (about 50%) of *Shh*-CKO;*Bax*-CKO hindlimbs. In contrast, in null *Shh*^-/-^;*Bax*-CKO limbs, *Alx4* and *Irx3* expression had already extended into the posterior limb bud by E10.75. Mutant numbers analyzed with the result shown are indicated in each panel. In remaining *Shh*-CKO;*Bax*-CKO embryos, expression was very similar to the null *Shh*^-/-^;*Bax*-CKO. Compass indicates limb bud orientation in all panels.

**Figure S4.**
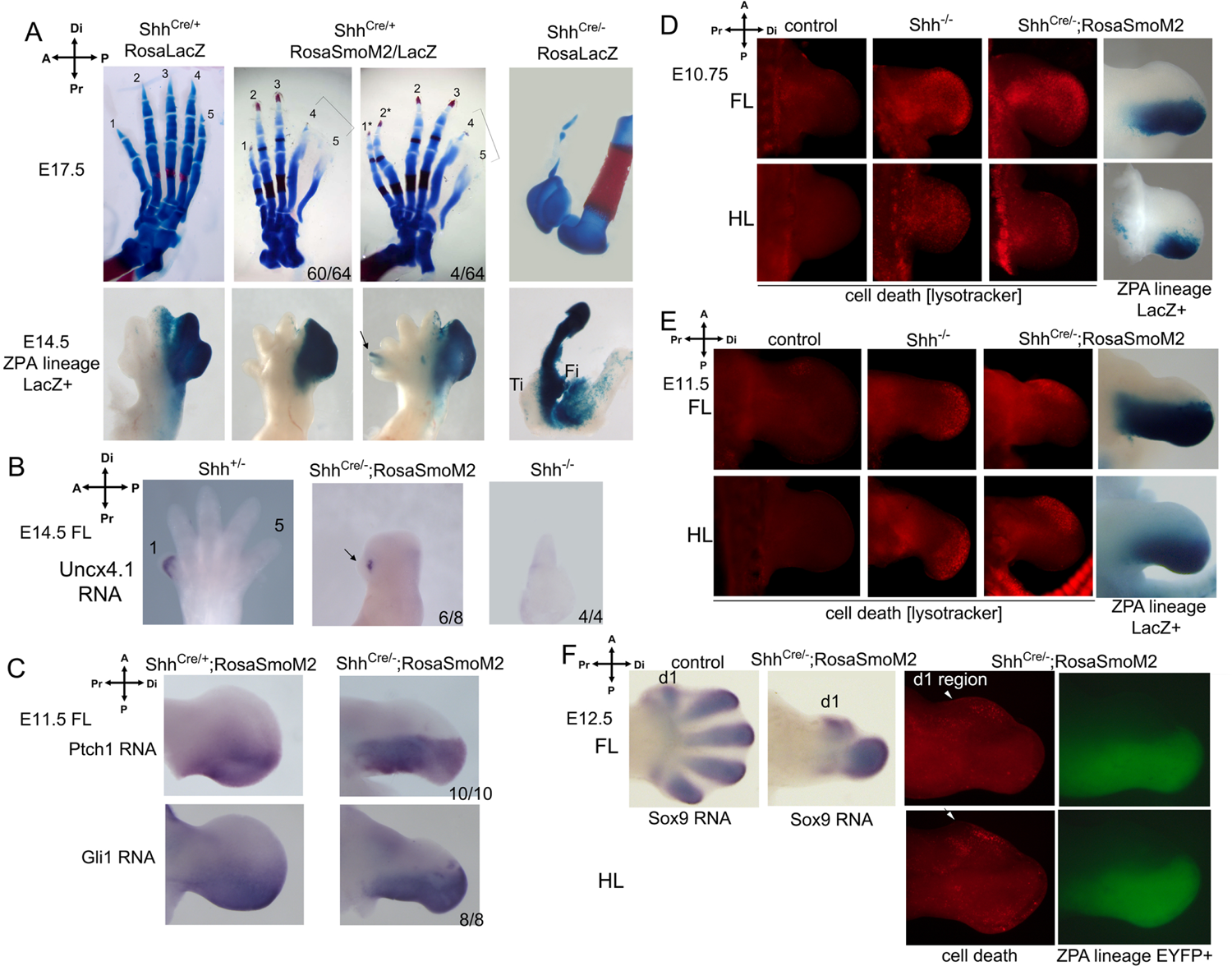
(related to Figure 6). A bona fide digit 1 is restored in *Shh*^cre/-^;Shh-SmoM2+ limbs and is not a consequence of cryptic anterior Hedgehog ligand/pathway activation. (A) Skeletal staining (E17.5) and ZPA lineage tracing (E14.5) of control *Shh*^cre/+^;Shh-SmoM2 (*Shh*+) embryos. In control *Shh*^cre/+^;Shh-SmoM2+, posterior digits are dysmorphic and uninterpretable because of constitutive Hedgehog pathway activation (brackets in middle 2 panels), and a small percentage of limbs have preaxial polydactyly (*, 4/64), related to ectopic anterior Shh activation revealed by ZPA lineage tracing at E14.5 (LacZ+, arrow). The single digit in *Shh* null arises entirely from ZPA descendants (right panels). Ti: tibia; Fi: fibula. Limb skeletons (and B panels) oriented with anterior (digit 1, tibia) at left, distal at top of panel. (B) *Uncx4.1* RNA expression in E14.5 forelimbs. *Uncx4.1* is expressed exclusively in digit 1 in control forelimbs (*Shh*^+/-^; left panel), and is expressed in the rescued digit 1 in *Shh*^cre/-^;Shh-SmoM2+ (6/8; middle panel, arrow), but is not detected in *Shh*^-/-^ limbs (n=4; right panel). For (C) - (F), all limb buds are oriented with anterior at top, distal at right of panel. (C) *Ptch1* and *Gli1* RNA expression in E11.5 forelimbs. *Ptch1* and *Gli1* are expressed in the posterior limb buds in *Shh*^cre/-^;Shh-SmoM2+, consistent with cell-autonomous Shh pathway activation in ZPA domain, but no expression is detected in the anterior limb bud (n=10, *Ptch1*; and n=8, *Gli1*). (D, E) Lysotracker staining (for cell death) and LacZ+ detection of ZPA lineage in *Shh*^cre/-^;Shh-SmoM2+ limb buds. Apoptosis persists in anterior non-ZPA descended limb bud in *Shh*^cre/-^;Shh-SmoM2+ limb buds similar to *Shh* null (*Shh*^-/-^) at E10.75 (D) and E11.5 (E). (F) Lysotracker staining (for cell death) and EYFP+ detection of ZPA lineage in *Shh*^cre/-^;Shh-SmoM2+ limb buds at E12.5 showing relationship of residual anterior apoptosis compared with early forming digit condensations visualized by Sox9 RNA in wildtype control and *Shh*^cre/-^;Shh-SmoM2+. Note that digit 1 condensation position at anterior-proximal limb bud border correlates with apoptosis negative zone in the mutant limb bud at same stage (white arrowheads). *Rosa*^LacZ^ or *Rosa*^EYFP^ Cre-reporters were used to mark ZPA lineage. Compass indicates limb bud orientation in all panels.

---

**Table S1.** Hindlimb phenotypes in sibling *Shh*-CKO embryos with or without *Bax* function for different Tamoxifen administration times (Hoxb6CreER activation).

| *Shh*-CKO;*Bak*<sup>-/-</sup>;Hoxb6CreER+ | 1 digit (KO) | 2 digits | 3 digits | 4 digits | 5 digits | % rescue |
| --- | --- | --- | --- | --- | --- | --- |
| **tamoxifen E9.5** |  |  |  |  |  |  |
| Bax[+]† | 12/12 | 0 | 0 | 0 | 0 | 0 % |
| Bax[-] | 16/16 | 0 | 0 | 0 | 0 | 0 % |
| **tamoxifen E9.5+3h (see also Fig 1)** |  |  |  |  |  |  |
| Bax[+] | 28/28 | 0 | 0 | 0 | 0 | 0 % |
| Bax[-] | 13/31 | 0 | 2/31 | 14/31 | 2/31 | 58 % |
| **tamoxifen E9.5+6h (E9.75)** |  |  |  |  |  |  |
| Bax[+] | 28/38 | 0 | 2/38 | 8/38 | 0 | 26 % |
| Bax[-] | 10/32 | 1/32 | 3/32 | 16/32 | 2/32 | 66 % |

†Bax[+] denotes *Bax*<sup>C/+</sup> genotype and Bax[-] denotes *Bax*<sup>C/C</sup> genotype, which becomes negative following Cre recombination.

